# Genomic profiling of cell lines reveals hidden research bias and caveats

**DOI:** 10.1101/2020.02.23.961516

**Authors:** Yongwen Luo, Lingao Ju, Yi Zhang, Yu Xiao, Xinghuan Wang

**Author notes:** **Corresponding author:** Dr. Xinghuan Wang.

## Abstract

Cancer cell lines are extremely valuable tools for carcinoma research. Biodiversity of cell lines and continuous random mutation in cell lines during passage, however, might result in phenotypic inconsistency and lead to biased experimental conclusions. Using statistics based on known and inferred protein interaction networks, as well as public research literature database, our study shows that essential driver genotypes of cell lines might have hidden impact on research results. Furthermore, by comprehensive genomic profiling of 8 most common used urothelial cell lines and comparing them to previous publications, we found that regardless of similar short tandem repeat (STR) profile, driver gene loss in cell lines could happen by random mutation. Our results suggest that clinical research using urothelial carcinoma cell lines might be influenced by cell line genotypes which could only be determined by next-generation sequencing. Meanwhile, this study indicates that the conditionally reprogrammed cells (CRCs), which closely resemble original tumor tissue, might represent a better alternative for in vitro research, which may be better used for personalized medicine.

---

Preexisting genetic mutations in cells might influence phenotypes caused by other molecular biology manipulations. For instance, the bladder cancer (BCa) cell RT4 harbors a *TACC3-FGFR3* fusion but not *TP53* mutation (Fig. 1F), while the other BCa cell T24 have *TP53* mutation but not *FGFR3* mutation[1]. To study whether these preexisting genotype could influence research conclusions, we performed a systematic review of the literature for urothelial carcinoma-related 1,589 articles either used or not used RT4 (Table S1). Firstly, we found significant association between the pathway which the target gene belongs and RT4 usage frequency: if the gene was known to associate with TP53 pathways, the research was less likely to include RT4 (Fig. 1A-B and Table S2).

**Figure 1.**
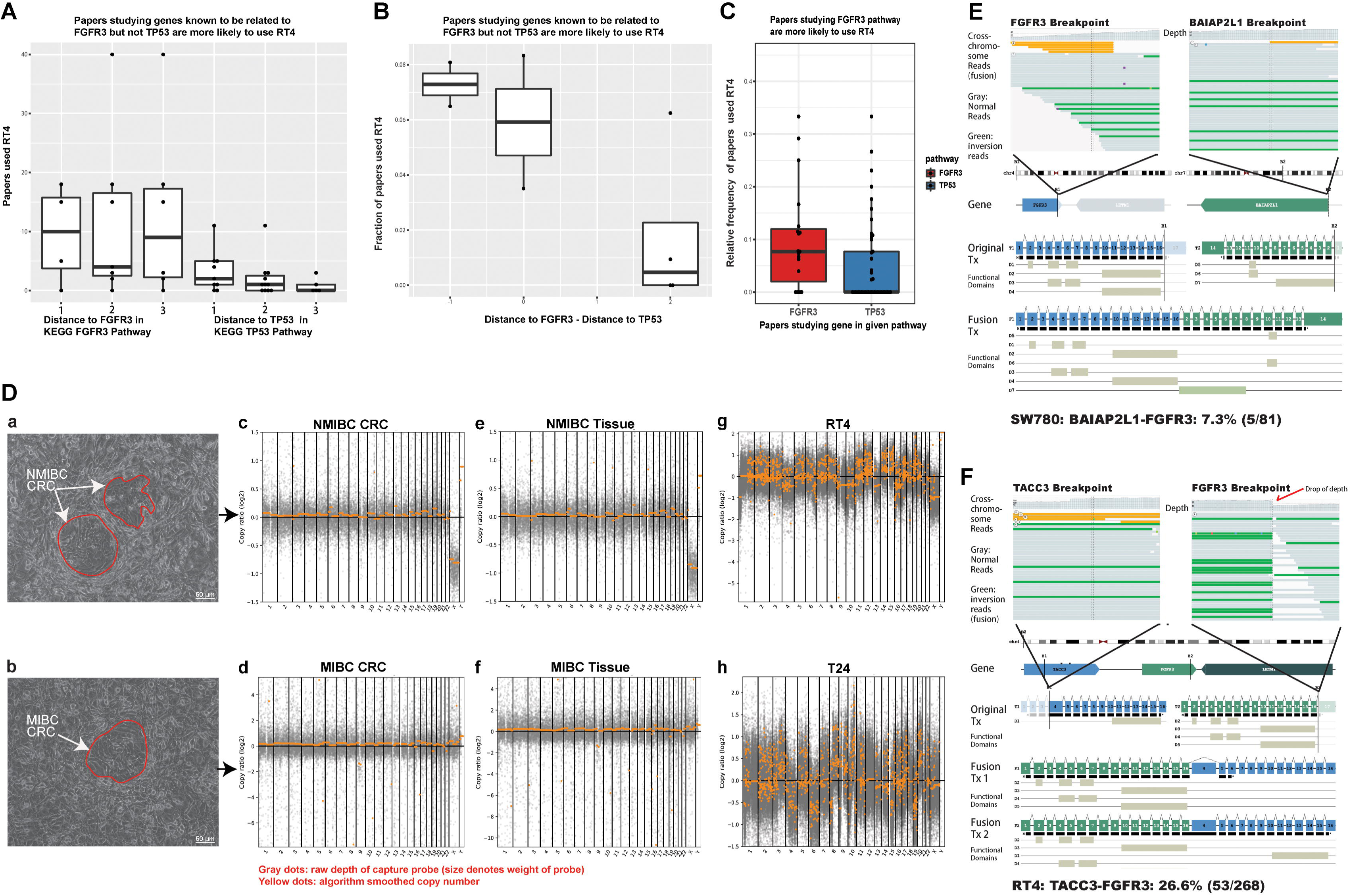
Comprehensive genomic profiling of Urothelial Carcinoma cell lines reveals hidden research bias and caveats. **(A)** Distance from given target gene to TP53 on KEGG pathway is negatively correlated with RT4 usage. X axis: distance to TP53 and FGFR3 (1, 2, 3 denotes direct interacting with TP53, one-step-away from TP53, and two-steps away from TP53, etc.). Y axis: PubMed search result number of target gene name with RT4. **(B)** The relative distance from given target gene to TP53 and FGFR3 based on protein-protein interaction network is correlated with RT4 usage. X axis: the difference between target-gene-to-TP53 distance and target-gene-to-FGFR3 distance. A minus number denotes the gene is closer to FGFR3 compared to TP53, and a positive number denotes the gene is closer to TP53 compared to FGFR3. Y axis: Frequency of RT4-containing search result in all PubMed search results with target gene and “bladder cancer cell line”. **(C)** Papers studying FGFR3 pathway are more likely to use RT4 density distribution of the RT4-containing search result frequency on PubMed for target genes in the KEGG FGFR3-related signalling pathway or TP53-related signalling pathway. **(D)** Morphology of 2 BCa patient-derived CRCs (a-b). Scale bar is 50 μm. Visually pan-genome copy number profile (yellow) with sequencing depth of probe region (gray) generated by CNVkit from 2 CRCs (c-d) and 2 BCa tissue (e-f) together with BCa cell lines T24 and RT4 (g-h) showing the significant deviation from genuine BCa cell lines. **(E)** Sequencing evidence for heterogeneity of driver oncogene fusion in SW780 cell line. Top panel showing the raw sequencing reads (yellow: cross-chromosomal DNA fragments; gray: normal DNA fragments; green: fragments in inverted direction) of FGFR3 (top-left) and BAIAP2L1 (top-right) loci. Middle panel showing the chromosomal bands and gene structure, denoting the breakpoint locus. Bottom panel showing the original transcript (Tx) and respective functional domains of each fusion partner gene, and the predicted fusion product transcript with a functional FGFR3 kinase domain and longer C-termini from BAIAP2L1. DNA fragments in support of fusion consists 7.3% (5 in 81) of all sequenced fragments, suggesting that the fusion driver oncogene is lost in a part of cells. **(F)** Sequencing evidence for *TACC3-FGFR3* driver oncogene fusion in RT4 cell line. Panel layout is similar to **(E)**. DNA fragments in support of fusion consists 26.6% (53 in 268) of all sequenced fragments, suggesting that the fusion driver oncogene likely to exist in homogeneous one-in-a-tetraploid cell state or in a heterogeneous manner. Copy number profile (a in panel D), however, did suggest that significant level of mosaicism exists in the RT4 population.

Then, we calculate the frequency of using RT4 and/or T24 for research on particular target gene (Table S3). Using PPI networks[2], we calculated the “interaction distance” between these gene to TP53 and FGFR3. Again, RT4 usage frequency is increased in research of genes with closer relationship to FGFR3 compared to TP53 (Fig. 1C). These results suggested the existence of latent bias in research: researcher might implicitly select consistent results by selecting carcinoma cell lines. The preexisting carcinoma cell line mutation landscape could determine its response to particular molecular biology manipulation, such response could be inconsistent. However, only consistent results would be reported.

Moreover, since genotype determined phenotype, identifying the genotype of carcinoma cell line is critical to the research result. State-of-the-art practices require genotyping cell lines using short tandem repeat (STR)[3]. However, STR only identifies whether cell lines are cross-contaminated, but does not fully report the genomic changes of cell lines. We collected 8 most widely used BCa cell lines and confirmed their identity by STR. Whole-exome sequencing (WES) of the 8 cell lines together with 2 conditionally reprogrammed cells (CRCs)[4] and 2 BCa tissues (Table S4) showed the CRCs have good consistency with tumor tissues, however, the cell lines showed significant deviation from genuine BCa by wide-spread shattering copy number variation (Fig. 1D and Fig. S1). Furthermore, comparing with the previous reports[5], we detected only a tiny fraction of the known driver FGFR3 fusion in SW780 (Fig. 1E). These results indicated although the carcinoma cell lines were not contaminated, they undergone pervasive genetic drift characterized by copy number variation and random loss of driver mutation.

Together, carcinoma cell line genotypes could influence research results by modifying molecular manipulation outcomes. Such influences could be implicit or explicit, and represents researcher’s selection bias in research. Furthermore, the genotypes undergone neutral genetic drift, which might lead to loss of important driver mutation and gain of novel genetic identity. Hence, our opioion suggests WES, instead of STR, should be widely used for determining carcinoma cell line genotype in future research. Meanwhile, the CRCs, a leading technology for living biobank, which closely resembles original tumor tissue, might represent a better alternative for *in vitro* research, which may be the direction of future precise medicine.

## Methods

### Literature search strategy and selection of studies

We performed an electronic search of PubMed (1966 to January 2019, *https://www.ncbi.nlm.nih.gov/pubmed*), for evaluating the relationship of preexisting genotype of BCa cell and research results on other genes. The search keywords were used with different combinations with both medical subject headings terms and text words: “(RT4[TW] OR RT-4[TW] OR T24[TW]) AND (“urinary bladder neoplasms” [MeSH Terms] OR “bladder neoplasms” [TIAB] OR “bladder cancer” [TIAB] OR “bladder tumour” [TIAB] OR “bladder tumor” [TIAB])”. Publication date was not restricted in our search. Reference lists of the included studies and supplemental materials were checked manually to further identify related studies. Three reviewers independently screened the title, abstract and keywords of each article retrieved. Full-text papers were screened for further assessment if the information given suggested that the study fulfilled the inclusion criteria and did not meet the exclusion criteria. Discrepancies were settled by discussion and consensus with all the authors.

### DNA extraction and NGS library preparation

Genomic DNA is extracted from cultured cells using Qiagen tissue DNA column. We processed the DNA according to a modified single-stranded DNA sequencing library prep protocol. Briefly, genomic DNA were sonicated, 3’-poly-A tailed with terminal transferase, ligated with a poly-dT-tailed P5 sequencing adaptor, and undergone several cycles of linear amplification using P5 sequencing primer. A P7 adaptor with random nucleotide 3’ overhang was then ligated to the 3’ end of linear amplification product. From there, sequencing library were amplified using P7+P5 primers to sufficient amount. Hybrid capture were performed using standard DNA probe capture practice with IDT XGEN whole exome panel. The post-capture libraries were sequenced on Illumina Novaseq.

### Cell culture

All urothelial cell lines were purchased from Cell Bank of the Chinese Academy of Sciences. The cell lines are authenticated via short tandem repeat (STR) analysis. The medium and culture conditions are as follows:

| Cell line | Medium supplemented with 10% FBS | Culture conditions |
| --- | --- | --- |
| RT4 | McCoy's 5A | 37°C, 5% CO<sub>2</sub> |
| T24 | RPMI-1640 | 37°C, 5% CO<sub>2</sub> |
| UM-UC-3 | MEM | 37°C, 5% CO<sub>2</sub> |
| J82 | MEM | 37°C, 5% CO<sub>2</sub> |
| SCaBER | RPMI-1640 | 37°C, 5% CO<sub>2</sub> |
| 5637 | RPMI-1640 | 37°C, 5% CO<sub>2</sub> |
| SW780 | L-15 | 37°C, free gas exchange with atmospheric air |
| SV-HUC-1 | F-12K | 37°C, 5% CO<sub>2</sub> |

### Bioinformatic pre-processing and determination of somatic mutation events

For SNV/indel and CNV calls, reads were trimmed using cutadapt to remove poly-T-tails, adaptors and the first 10bp from each end before being mapped to GRCh37+decoy reference genome using bwa-mem. For structural variation calls, reads were only trimmed to remove adaptors before mapping. Before calling any variants, the bam files were duplication-marked, realigned, and passed through a BQSR pipeline. SNV (including single nucleotide variation and small indel, hereinafter designated as SNV) calls are made with Sentieon TNscope with sequenced NA12878 cell line as control, or with Pisces with tumor sample only. The tumor cell and control samples additionally run through a Sentieon Haplotyper pipeline individually to call AF > 0.2 variants. The TNscope callset were filtered with AF(tumor) > 0.2, ALT_F1R2 > 5, ALT_F2R1 > 5, ALT >= 30, AF(tumor)/AF(control) > 5, AF(normal) < 0.01. The Pisces callset were used to calibrate oxo-G levels using VQR. After calibration, Pisces callset were intersected with TNscope callset to generate a “somatic callset”. Finally, germline mutations from the normal sample Haplotyper callset were removed from the somatic callset. Annotations were done with Annovar and VEP. We further annotated the mutations with gnomAD *(http://gnomad-old.broadinstitute.org/)*, MCAP, Spidex, SCADA, Clinvar *(https://www.ncbi.nlm.nih.gov/clinvar/)* and public HGMD databases (*http://www.hgmd.cf.ac.uk/ac/index.php*). Sequenza was used to process tumor sample to estimate cellularity (tumor fraction). To call somatic CNV, copy-number-homogeneous segments were generated using CNVkit run with tumor and paired normal samples, and filtered with cellularity. We passed BAF from the somatic callset as well as germline Haplotypecaller callset into sequenza and CNVkit, to estimate correct copy number information for each segment. Structural variation was called and annotated using the iCallSV pipeline. Mavis (using Lumpy) was also used to search for possible low-frequency SV. MSI score is calculated with MSIsensor. To calculate mutation signature, the filtered, passed, main- and subclonal mutations were passed to deconstructSig.

### Quality control of sequenced samples

Quality control of the files was done with in-house script. We checked the mapping quality, mapping rate, duplication rate and uniformity to control for molecular assay and sequencing. Tumor samples in this study were sequenced to a median of > 200x. Failed regions (< 5x) were < 0.2% in any sequenced samples. For each tumor sample, > 98% of captured regions achieved a minimum of 50x.

Detailed information is shown in the Supplementary information.

## Abbreviations

BCa: bladder cancer
CRCs: conditionally reprogrammed cells
STR: short tandem repeat
WES: whole-exome sequencing

## Declarations

### Ethics approval and consent to participate

The study using clinical information and human samples (including surgical tissue specimens and primary cancer cells) was approved by the Ethics Review Committee at Zhongnan Hospital of Wuhan University (approval number: 2015029). Human sample preservation by the Zhongnan Hospital Biobank, the official member of the International Society for Biological and Environmental Repositories (https://irlocator.isber.org/details/60), was approved by the Ethics Review Committee at Zhongnan Hospital of Wuhan University (approval number: 2017038) and China Human Genetic Resources Management Office, Ministry of Science and Technology of China (approval number: 20171793).

### Consent for publication

All authors give consent for the publication of manuscript in Molecular Cancer.

### Availability of supporting data

Detailed information is shown in the Supplementary information. The datasets used and/or analysed during the current study are available from the corresponding author on reasonable request.

### Authors’ contributions

YL and LJ performed experiments. All authors contributed to the research. YL, LJ, YZ, YX and XW designed the study and wrote the manuscript. All authors approved the final manuscript.

## Acknowledgments

We thank the patients and their family members for participating in our study. We gratefully acknowledge excellent technical assistance provided by Ms. Yuan Zhu, Ms. Mengxue Yu and Ms. Yayun Fang from Zhongnan Hospital of Wuhan University.

