## Supplementary information for "Genomic profiling of cell lines reveals hidden research bias and caveats"

**Supplementary material and methods**

**Patient selection and pathological classification**

The clinical, pathological, follow-up data records of two bladder cancer (BCa) patients were collected (Table S4), and two pathologists were invited to confirm the histology diagnosis independently. The study using clinical information and human samples (including surgical tissue specimens and primary cancer cells) was approved by the Ethics Committee at Zhongnan Hospital of Wuhan University (approval number: 2015029). The human sample preservation by Department of Biological Repositories at Zhongnan Hospital of Wuhan University, the official member of the International Society for Biological and Environmental Repositories-International Repository Locator (ISBER-IRL)[1], was approved by the Ethics Committee (approval number: 2017038) and China Human Genetic Resources Management Office, Ministry of Science and Technology of The People’s Republic of China (approval number: 20171793). All BCa patients provided the written informed consent. The procedures in this study were done in accordance with the ethical standards of the institutional ethics review committee.

**Conditional reprogrammed cells (CRC) technology**

The method used to establish patient-derived conditional reprogramming (CR) bladder cancer cell model from fresh BCa tissues was performed according to Liu *et al.* with minor modifications[2]. Briefly, two fresh BCa tissues were rinsed with 95% ethanol (up to 3 seconds) and cold PBS. Chop the samples into small (< 10 mm) pieces with scissors. Then transfer the tissue fragments to 4 ml F medium, containing collagenase, hyaluronidase and dispase, and incubate for 2 hours at 37 °C on a rocking platform. After dissociation, the cell pellet was collected by centrifuging for 5 minutes at 500 g and resuspend in 10 ml complete DMEM. The cell suspension was filtered by 100 mm cell strainer, followed by centrifugation for 5 minutes at 300 g. Then plate the processed cell pellet in a T25 flask together with Swiss-3T3-J2 mouse fibroblasts feeder cells in complete F medium. Replace feeder cells every three days and observe colonies of CR prostate cancer cells by phase contrast microscopy (Leica Ltd., Germany).

**Determine driver variants**

Mutations were classified using gnomAD prevalence, pathogenicity prediction, and known evidence from Clinvar and HGMD open. For LP and PAT mutations in HGMD, we used a population frequency cutoff of 0.002. For gnomAD alleles, we used a population frequency cutoff of 0.001, East Asian allele count (AC_EAS) < 10, and homozygous East Asian donor count (Hom_EAS) < 2. Alleles without gnomAD frequency were annotated against an in-house population frequency library consisting 5000 Chinese WES/WGS data. We filtered severe mutations (frameshift, inframe-del, inframe-ins, stop-gain, splicing defect with SCADA > 0.8 or abs(Spidex) > 3). For missense or indel mutations, we require the locus MCAP > 0.02. For common known oncogene and tumor suppressors, we further validated mutations using the list provided by Bailey et al 2018 and Clinvar[3, 4]. Identification of missense driver mutations is done by filtering against COSMIC database (*https://cancer.sanger.ac.uk/cosmic/signatures/ID*).

To calculate the cancer specificity of any missense driver event, we computed the relative prevalence of the mutation in gnomAD plus the in-house Chinese germline database and compared it to the prevalence of this mutation in TCGA (*https://portal.gdc.cancer.gov/*) and COSMIC database using Fisher’s exact test. We found that a prevalence of 5 in COSMIC v.88 database, or 3 in TCGA MC3 MAF file, with a gnomAD global population frequency lower than 0.1%, is adequate to determine that the mutation is recurrent and somatic in nature for tumor tissues. For LOF-driver genes, we also included likely pathogenic splicing/start-lost/stop-gain/frameshift mutations, as well as CNV-loss. For GOF-driver genes, we included high level CNV-gain (normalized to >= 5x). Structural variation results were filtered to include only the kinase-domain-included, activating translocations, and a frequency of > 25%.

**Supplementary Figures**

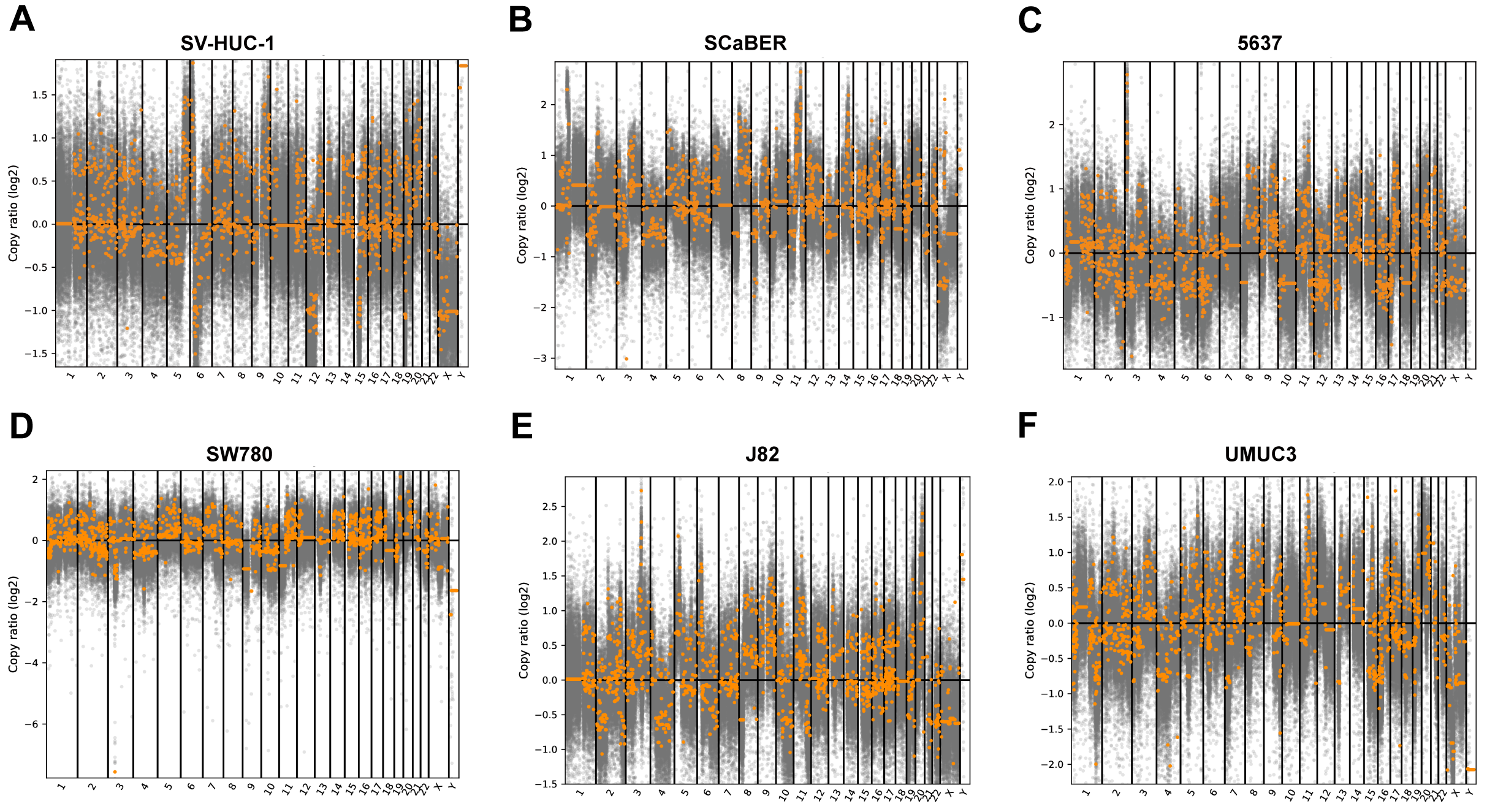

**Figure S1. Pan-genome copy number variations plots of urothelial cells.** Visually pan-genome copy number profile (yellow) with sequencing depth of probe region (gray) generated by CNVkit from SV-HUC-1, SCaBER, 5637, SW780, J82 and UMUC3.
